# Phage-plasmids spread antibiotic resistance genes through infection and lysogenic conversion

**DOI:** 10.1101/2022.06.24.497495

**Authors:** Eugen Pfeifer, Rémy A. Bonnin, Eduardo P.C. Rocha

## Abstract

Antibiotic resistance is rapidly spreading by horizontal transfer of resistance genes in mobile genetic elements. While plasmids are key drivers of this process, very few integrative phages encode antibiotic resistance genes. Here, we find that phage-plasmids, elements that are both phages and plasmids, often carry antibiotic resistance genes. We found 60 phage-plasmids with 184 antibiotic resistance genes, including broad-spectrum-cephalosporins, carbapenems, aminoglycosides, fluoroquinolones and colistin. These genes are in a few hotspots, seem to have been co-translocated with transposable elements, and are often in class I integrons, which had not been previously found in phages. We tried to induce six phage-plasmids with resistance genes (including four with resistance integrons) and succeeded in five cases. Other phage-plasmids and integrative prophages were co-induced in these experiments. As a proof of principle, we focused on a P1-like element encoding an extended spectrum β-lactamase, *bla*_CTX-M-55_. After induction, we confirmed that it’s capable to infect and convert four other *E. coli* strains. Its re-induction led to further conversion of a sensitive strain, confirming it’s a fully functional phage. This study shows that phage-plasmids carry a large diversity of clinically relevant antibiotic resistance genes that they transfer across bacteria. As plasmids, these elements seem very plastic and capable of acquiring genes from other plasmids. As phages, they may provide novel paths of transfer for resistance genes, because they can infect bacteria distant in time and space from the original host. As a matter of alarm, they may also eventually mediate transfer to other types of phages.

**Importance:** Dissemination of antimicrobial resistances is a major threat to global health. Here, we show that a group of temperate bacterial viruses (=phages), termed phage-plasmids, commonly encode different and multiple types of resistance genes of high clinical importance, often in integrons. This is unexpected since phages typically do not carry resistance genes and, hence, do not confer their hosts with resistance upon infection and genome integration. Our experiments with phage-plasmids isolated from clinical settings confirmed they infect sensitive strains, rendering them antibiotic resistant. The spread of antibiotic resistance genes by phage-plasmids is worrisome because it dispenses cell-to-cell contact, necessary for the canonical plasmid transfer (=conjugation). Furthermore, their integrons are now genetic platforms for the acquisition of novel resistance genes.

## Introduction

Antimicrobial resistances (AMR) are fast disseminating among human-associated bacteria and have been classified as major challenges to Global Health (1). Enterobacterales are identified as the most critical group (2) against which new drugs need to be developed. Resistance is the result of one of multiple mechanisms: limiting drug uptake; target modification; active drug efflux and drug inactivation. The latter includes extended spectrum ß-lactamases (e.g. ESBLs) that allow Enterobacterales to become resistant against most ß-lactams (such as penicillins or broad-spectrum cephalosporins). Although ESBLs do not provide directly resistance to carbapenems (last-resort antibiotics within ß-lactams), the wide and improper use of carbapenems, especially as a first-line treatment, has promoted the emergence of carbapenem-resistant Enterobacterales (CRE) strains that are commonly found to be resistant to others antibiotic classes (3). While low-level resistance to ß-lactams can be provided by many mechanisms such as qualitative or quantitative modifications of porins, high resistance is usually associated with the acquisition of genes encoding ESBLs or carbapenemases by horizontal gene transfer (4). The most important and clinically relevant carbapenemases identified in Enterobacterales belong to class A (KPC-like enzyme), class B (NDM-, VIM-and IMP-like enzyme) and class D (OXA-48-like enzyme) type ß-lactamases (5). Plasmids are key drivers of the transmission of antibiotic resistance genes (ARGs) between bacteria, usually by conjugation (6, 7). Transfer is also facilitated by the presence of mobile genetic elements (MGEs) translocating genetic information between replicons (8). Notably, ARGs are often flanked by transposable elements that facilitate their translocation between plasmids or between plasmids and the chromosome (9). Integrons can also facilitate the translocation of ARG cassettes (10). Mobile integrons are usually associated with plasmids and/or transposons and consist of one integrase (here of the type IntI1) and a small array of gene cassettes flanked by recombination sites. Integrons can acquire new gene cassettes from other integrons and shuffle the existing ones (11). A large fraction of the cassettes of mobile integrons consists of ARGs (10). The co-transfer of multiple ARGs in an integron facilitates the emergence of multi-drug resistance strains.

Temperate bacteriophages (phages) can mobilize genes by different types of transduction processes (generalized, specialized and lateral) (12) or introduce new genes by lysogenic conversion (13). Generalized transduction relies on erroneous packaging of non-phage DNA by specific types of phages and tends to occur at low frequencies (14), whereas lateral and specialized transduction require proximity between the transferring genes and the phage (12). All these processes have been shown to result in the transfer of ARGs in the lab, but there is extensive controversy on the extent and pertinence of this process in natural environments (15–19). In contrast, lysogeny is common in nature (20–22). In this case, the phage remains mostly silent in the cell (as prophage), but accessory genes can be expressed and change the host phenotype. Many toxins with key impact on the virulence of bacterial pathogens are present and expressed from prophages (13). However, very few phages encode *bona fide* ARGs (16). To the best of our knowledge, no natural phage with ARGs has been shown to be fully functional – i.e., to lyse the original host cell, infect another cell and then repeat the cycle to infect a third cell – and provide antibiotic resistance by lysogenic conversion.

While most prophages integrate the chromosome, some remain in cells as phage-plasmids (P-Ps). These are temperate phages that transfer horizontally (infect) as viruses but remain and replicate within cells as plasmids. In a previous work, we found P-Ps to be numerous, widespread and organized in different groups (23). A few of these groups are frequent in enterobacteria and other important nosocomial pathogens, *e*.*g*. P1-like P-Ps are very frequent in *Escherichia coli*, SSU5-like and N15-like elements in *Klebsiella pneumoniae*, and AB-like P-Ps in *Acinetobacter baumannii*. P-Ps tend to be larger than prophages integrated in the chromosome. The P-Ps have loci that are very plastic and contain genes typical of plasmids and other more conserved loci encoding phage-related genes (23). Some of the P-Ps, notably the P1-like, can also be efficient transducers (24). The double nature of P-Ps, being a plasmid and a phage, led us to think that they might contribute more, especially by lysogenic conversion, to the spread of ARGs than the other phages. Furthermore, a few reports have identified elements resembling P-Ps carrying ARGs. For example, P1-like elements were identified encoding an *mcr-1* gene conferring resistance to colistin in *K. pneumoniae*, and ESBLs in *Salmonella* spp. and *E. coli* but induction and transmission could not be confirmed (25–27). Recently, a P1-like element with several predicted ARGs could lysogenize one commensal *E. coli* strain and provide resistance to streptomycin (28). This shows that P-Ps can carry and transfer ARGs, although the viability of the full phage lifecycle (infection and reinfection) was not yet confirmed.

Here, we test the hypothesis that P-Ps are more likely to encode ARGs than the other phages because they share characteristics of plasmids such as presence of transposable elements and regions of high genetic plasticity. For this, we searched a large number of P-Ps, plasmids and phages from reference databases for *bona fide* ARGs. We found many ARGs and their acquisition seems to have been driven by transposable elements and integrons. To test if the P-Ps can be induced we scanned a collection of carbapenem-resistant strains for putative P-Ps. The tested cases showed almost systematic induction of P-Ps. Among those induced, we then tested if P-Ps were able to convert a panel of sensitive strains into bacteria resistant to broad-spectrum cephalosporins.

## Methods

### Genomic data

We used the completely assembled genomes of 8399 bacterial strains, including their 21550 plasmids, and the completely assembled genomes of 3725 phages. All genome data was retrieved from the non-redundant NCBI RefSeq database (29) (March, 2021).

### Similarity between mobile genetic elements

The weighted gene repertoire relatedness (wGRR) assesses the similarity of gene repertoires between pairs of mobile genetic elements, by taking into account their number of bi-directional best hits (BBH) and their sequence identity. It is computed as described previously (23) for all genomes/contigs of phages, plasmids and P-Ps. Briefly, MMseqs2 (v. 13-45111) (30) was used to conduct an all-vs-all gene comparisons between the elements. BBHs between two genomes were extracted if they met the following criteria: evalue <10^−4^ and sequence identity >35% covering at least 50% of both gene sequences. wGRR was computed as:

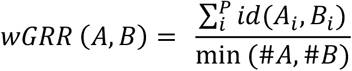

A_i_ and B_i_ are the *i*th BBH pair of *P* total pairs. The gene number of the smaller genome is min(#A,#B), and the sequence identity between the BBH pair is id(A_i_,B_i_). The sum of the sequence identities (of the BBHs) normalized to the gene number of the smaller genome is defined as the wGRR between the two genomes.

### Identification, and classification of phage-plasmids (P-Ps)

P-P genomes were identified as described previously (23). Briefly, we searched for genes encoding phage-like functions in plasmids of intermediate size (>10kb and <300 kb) by using carefully-selected pVOG (31), PFAM (32) and TIGRFAM (33) HMM protein profiles. The detection used HMMER v 3.3.2 (34). A positive hit was assigned if the alignment covered at least 50% of the protein profile with a domain i-Evalue <10^−3^. The distributions of hits in the plasmids were given to previously trained random forest models that provided the list of putative P-Ps. dsDNA Phages (larger than 10 kb) were screened for plasmid functions using protein profiles specific for plasmid replication and partition systems (35). Phages with hits for plasmid functions were extracted and were compared with plasmids and P-Ps (23). Novel elements having wGRR >= 0.4 with elements present in the list of previously identified P-Ps were added to the list of putative P-Ps. This resulted in 1416 putative P-Ps, including 740 previously identified.

The classification of novel P-Ps is based on the similarity to previously identified P-P groups. P-Ps that were not identified in our previous study (23), typically because they correspond to more recent genome sequences, were assigned to defined P-Ps groups when they have wGRR >= 0.5 and at least half of their genes homologous to a previously classed P-P. When there are multiple hits, the P-P was classed according to the classification of the element with the highest wGRR.

### Identification of antibiotic resistance genes (ARGs), IS elements and integrons

We searched genomes for ARGs using as a reference the databases CARD (36), ResFinder (37), and ARG-ANNOT (38). We searched for sequence similarity between the genes of a MGE (phage, plasmid, P-P) and these databases using blastp (v.2.12.0+) (39) (to compare with protein sequences of CARD and ARG-ANNOT) and blastx (v.2.12.0+) (39) (for nucleotide sequences of ResFinder). We collected all hits in all databases respecting the following constraints: evalue <10^−5^, sequence identity ≥99% and alignment covering sequences by ≥99%. The results were compared with the output of AMRFinderPlus (3.10.18) (tool from NCBI for ARG detection (40)) (supplementary figure S2). IS elements were identified using ISEScan (v. 1.7.2.3, default parameters) (41). Integrons were identified using IntegronFinder (v. 2.0rc6, default parameters) (42).

### Pangenome graphs

To compute pangenomes of P-P groups (including newly assigned members), we followed the same workflow as described previously (23). We computed the pangenome with PPanGGolin (v. 1.1.136, default parameters) (43). Genes (including ARGs) were grouped into gene families, if they had an identity of at least 80% covering 80% of the sequence. We made the visualization of the pangenome graphs with Gephi (https://gephi.org/) and igraph (https://igraph.org/r/) in the R environment.

### ARG-encoding P-Ps in carbapenem-resistant *Enterobacteriaceae*

Draft genomes of carbapenem-resistant Enterobacterales (CRE) received from the French National Reference center were screened for ARG-containing P-Ps. For this, we predicted the genes using prodigal (v2.6.3, with default parameters) (44) and compared each contig with known P-Ps using the wGRR. We selected contigs with wGRR >= 0.4 for further study. These contigs were annotated in terms of ARGs using the same method as that used for the P-Ps (see section on the identification of ARGs).

Strains with contigs that were regarded as parts of putative P-Ps and encoding ARGs, were then re-sequenced using long reads. Cells were cultivated in 4 ml LB-Miller medium (w/ Carbenicillin 50 µg/ml at 37 **°**C, 250 RPM) for ∼16h, pelleted and their DNA were isolated with a modified version of the guanidium thiocyanate method (prior to DNA precipitation samples were treated with RNase A at 37 **°**C for 30 min) (45). DNA library preparation (SMRTBell Library 10 kb insert size) and sequencing was done with the Biomics sequencing platform of the Institut Pasteur (C2RT) (Paris, France) with the technology of Pacific Biosciences. The obtained reads were assembled by flye (v.2.7.1-b1590) (46) with default parameters (see supplementary figure S8).

### Growth experiments with Mitomycin C (MMC)

The CRE strains with ARG-encoding P-Ps were first cultivated in 4 ml LB-Miller medium w/ Carbenicillin 50 µg/ml (37 **°**C, ca. 16h). The stationary cultures were diluted 1:100 and cultivated in a 96-well plate (200 µl per well) in LB-Miller medium w/ Carbenicillin 50 µg/ml for 1h. Subsequently, Mitomycin C (MMC) (Sigma-Aldrich, St. Louis, United States) was added in final concentrations of 5 µg/ml, 1 µg/ml and w/o MMC. The growth was monitored by following the absorbance at OD_600_ measured with a TECAN Genios™ plate reader (Månnedorf, Switzerland) (supplementary figure S9).

### Polyethylene glycol (PEG) precipitation of phage virions

CRE strains w/ P-Ps were cultivated as described in the MMC growth experiment. 4 ml-cultures were started in LB-Miller w/ Carbenicillin 50 µg/ml by using 1:100 dilutions of the overnight cultured strains and 5 µg/ml MMC was added after 1h. 4h after the MMC addition, samples (2ml) were taken for PEG precipitation, pelleted and the supernatant was filtered (0.22 µm) (=phage lysate). To these phage lysates, 5xPEG/NaCl (PEG-8000 20%, NaCl 2.5 M) solution was added in a 1:5 ratio, inverted several times and chilled on ice for 1h. Subsequently, virions were pelleted at 3 min and 13 000 *g* and the supernatant carefully discarded. The pellets were resolved in TBS buffer (50 mM Tris-HCl pH 7.5, 150 mM NaCl) in 1/10 of the initial phage lysate volume and incubated for another hour on ice. The PEG precipitated samples were further used for phage DNA extraction or infection experiments.

### Extraction and sequencing of DNA located in virions

Virion DNA was extracted as described by Jakočiūnė and Moodley (47) after PEG precipitation and starting from step 3.2. Residual bacterial DNA was removed by treating samples with DNase I and RNase at 37 **°**C. The phage protein capsid was digested with Proteinase K and the DNeasy Blood & Tissue Kit (Qiagen, Hilden, Germany) was used to purify the DNA. Quantity and quality of purified DNA was checked by a Qubit™ fluorometer and a NanoDrop™ spectrometer. Library preparation (Illumina^®^ TruSeq™ DNA PCR-Free), sequencing and quality checks were done by the Biomics sequencing platform of the Institut Pasteur (C2RT) (Paris, France) by short-reads (paired-end, 250 bp length) on a MiSeq system (Illumina^®^, San Diego, U.S.).

### Sequence data processing

We took DNA obtained after the MMC induction experiment and tried to assemble the P-Ps. However, given the presence of repeats in these elements, they were not fully assembled. To obtain the complete sequences, we put together the long reads from the genome sequencing (obtained before) and the short reads from the MMC induction experiment (see section above). These were then co-assembled using Unicycler (v. 0.4.8) (48) with default parameters. The hybrid assembly resulted in 4-15 linear and/or circular contigs per strain representing the sequence of induced P-Ps, prophages and other DNA found in virions after MMC treatment (supplementary figure S10). We evaluated the assemblies by checking if the P-P contigs were closed (fully assembled) or if they weren’t, by comparing them with known P-Ps (supplementary figure S11). Subsequently, we mapped the reads (obtained after the MMC induction) on these assemblies to assess how they cover them using bowtie2 (v. 2.4.4) (49) with default parameters. To extract the coverage, we converted the output SAM-files to sorted BAM-file using SAMtools (v. 1.13) (50) and obtained the coverage with BEDTools (v2.30.0) (51). In addition, we computed the (background) coverage caused by undigested gDNA. For this, we took the short reads (from the MMC induction experiment) that did not map on the hybrid assembled contigs (= DNA outside of virions) and aligned them on the contigs acquired from the genome sequencing experiment. The mean coverage was computed by dividing the absolute read coverage per contig (genome) by the size of the contig (genome).

### Generation of antibiotic resistant phage-plasmid lysogens

PEG precipitated phage lysates were prepared and stored at 4 **°**C. Potential host strains were cultivated the day before in 4 ml LB-Miller medium for approx. 16h at 37 **°**C and 250 RPM. The stationary cultures were diluted 1:100 in LB-Miller medium and grown until an OD_600_ of 0.5 to 1. Subsequently, 50 µl of the phage lysate was added to 50 µl host culture w/ 2mM CaCl_2_ and incubated under non-shaking conditions at 37 **°**C for 1h. After incubation, the cell/phage-lysate mixture was plated on agar plates with the required antibiotic concentration (to screen for lysogens). Antimicrobial susceptibility tests were performed as described (52) and interpreted according to the EUCAST guidelines. Colonies were tested by PCR for the presence of P-P genomes (amplifying two regions; for region 1 with 522 bp PP-R1A 5’-CTACCAGACCGCCTTTCTCAAAC-3’, PP-1B: 5’-TTGCCGAAACTAGAGAATAAATACGG-3’ and for region 2 with 423 bp PP-R2A 5’-TTAACCTTTGTCGGCGTCGG-3’, PP-R2B 5’-ATGTCATTCTTTTCTACATTAAAAACAGC-3’) and finally confirmed by genome re-sequencing. Genomic DNA was isolated as described in the section on ARG-encoding P-Ps in CRE strains. Library preparation (Illumina^®^ TruSeq™ DNA PCR-Free) and sequencing was done on the C2RT Biomics platform of the Institut Pasteur (using short reads, paired-end, 250 bp) on an Illumina MiSeq system. The resulting reads were mapped on the P-Ps and on the host genomes (*E. coli* 55989: NC_011748, *E. coli* CIP 105917: NZ_CP041623, *E. coli* CIP 53.126: NZ_CP022959, *E. coli* CIP 76.24: NZ_CP009072). Coverage was extracted as described in the section on processing sequencing data.

### Data processing, storage and availability

If not otherwise stated, all analysis and illustrations were done in the R environment (https://www.r-project.org/) with Rstudio (v. 1.4.1106). All reads were uploaded to the European Nucleotide Archive (ENA) (https://www.ebi.ac.uk/ena). Short and long reads from the MMC induction, the genome sequencing experiment, the verification of P-P acquisition as well as the P-P nucleotide sequences gained by the hybrid assemblies are accessible under the following ENA study number PRJEB52357. Details on accession numbers and experiments are listed in supplementary table S5.

## Results

### Antibiotic resistance genes are common in phage-plasmids but rare in other phages

To assess quantitively the distribution of ARGs in plasmids, phages, and P-Ps, we searched for these genes in the complete bacterial and phage genomes of the RefSeq database. For this, we first updated our database of P-Ps using a previously described detection method (in (23)) (supplementary figure S1A). The novel P-Ps were classed in groups using their similarities to previously classed elements as measured by the weighted gene repertoire relatedness (wGRR) (see methods and supplementary figure S1). This led to an almost doubling of the database of P-Ps to a total of 1416 P-Ps. These elements represent 5.6% of the 25,275 phages and plasmids.

We searched for genes in phages, plasmids and P-Ps with very high sequence similarity (at least 99% identity and 99% coverage) to verified ARGs from three reference databases (ARG-ANNOT, ResFinder and CARD). In agreement with previous studies (8), ARGs were frequently found in plasmids (20.8%) and almost never found in phages (2 out of 3585 genomes, <1 ‰) (see figure 1). A total of 4.2% of the P-Ps carried ARGs, a frequency that is intermediate between that of phages (ca. 76.0 times more) and plasmids (4.9 times less). To further validate the annotation of ARGs, we compared the results of the three databases with the analysis of our data using the NCBI AMRFinderPlus software (40). We found similar ARGs in P-Ps and phages, and an increase in the number of plasmids with ARG of about 13.5% (supplementary figure S2). In P-Ps, the ARGs encode a variety of enzymes e.g. β-lactamases, dihydrofolate reductases, and aminoglycosides-modifying enzymes. We also identified a few genes encoding efflux pumps (supplementary table S1). Overall, our analysis shows that P-Ps encode ARGs much more often than the remaining phages. In some cases, they encode resistance genes to last-line antibiotics, like the *mcr-1* against colistin, various *blaKPC* (type 2, 3, 4 and 33) and *blaNDM-1* genes against carbapenems (supplementary table S1).

**Figure 1:**
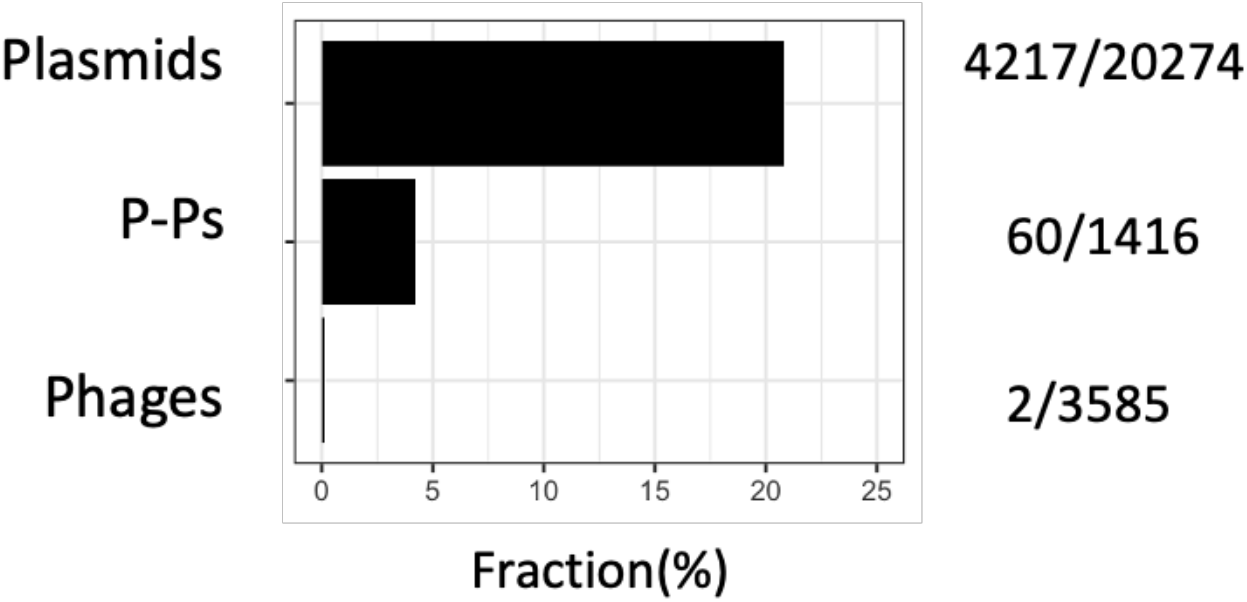
Number of mobile genetic elements encoding ARGs. The values after the bars indicate the number of elements encoding ARGs over the total number of elements considered in the analysis.

### Resistance genes are in specific types and loci of phage-plasmids

Most of the P-Ps carrying ARGs (47 of the 60 P-Ps) were found in genomes of just four species: *Acinetobacter baumannii* (n=8), *Escherichia coli* (n=20), *Klebsiella pneumoniae* (n=14) and *Salmonella (spp*. and *enterica)* (n=5). This is not overly surprising; our previous study showed these species had many P-Ps (23), many genomes of these species are available in the database, and these are all pathogenic bacteria known to develop antibiotic resistance (53). The majority of these P-Ps were assigned to well-defined P-P groups (23). P1-like P-Ps represent a third of the elements with ARGs (21 cases) of which all are in the P1-subgroup 1. We also detected 12 SSU5-related P-Ps and 8 AB-like P-Ps with ARGs (supplementary table S1). Interestingly, we could not detect any ARGs in P-Ps of the N15 group, the pMT1 group and the P1 subgroup 2. The results for N15 are particularly intriguing, because these elements are very abundant in nosocomial species, like *E. coli* and *K. pneumoniae* (23).

We analyzed the genomic locations of ARGs in P-Ps to shed light on how these genes were acquired and how these events may have affected the genetic organization of P-Ps. For this, we computed the pangenomes of the P-P groups, selected gene families present in high or intermediate frequency in the pangenomes to build a graph of the genetic organization of the elements, and placed the ARGs in relation to this backbone (figure 2). ARGs were never found in the persistent genome of the P-Ps, in light with the hypothesis that they were recently horizontally acquired and that they are not essential. Some P-Ps harbor one ARG, but the majority (n=39) has multiple genes, with up to 13 ARGs detected in a single putative P-P (pASP-135, NZ_CP016381) from the *Aeromonas hydrophila* strain AHNIH1 (supplementary table S1). This fits previous suggestions that *Aeromonas* spp have a key role in the genetic transfer of ARGs (54). One of the ARGs is the *bla*_KPC-2_ conferring resistance to carbapenems and is of great concern since it could act as a reservoir for this gene. Genes that commonly promote recombination and genomic plasticity, such as transposases and recombinases, were systematically identified in close proximity to the ARGs (supplementary figure S3-S7). Transposases of the IS*6*-like family were particularly frequently found next to the ARGs, especially those of the type IS*26* (figure 2). This family of ISs has been previously involved in the spread of clinically relevant ARGs, commonly causes plasmid co-integration, and its insertion results in hybrid promoters that can influence the expression of neighboring genes (55). Notably, most ARG families (21/24) of the P1 subgroup 1 are close to an IS*6*-like transposase (figure 2), suggesting that this transposable element drives the ARG acquisition in these P-Ps. In addition, in the AB, pKpn, and SSU5_pHCM2 groups some ARGs are found next to IS*5*-like, IS*30*-like, Tn*3*-like transposases, and several other types of recombinases. In the pSLy3-like group, we found no IS6-like transposases next to ARGs, but we did find an IS*Ec9* transposase (figure 2, supplementary figure S7).

**Figure 2:**
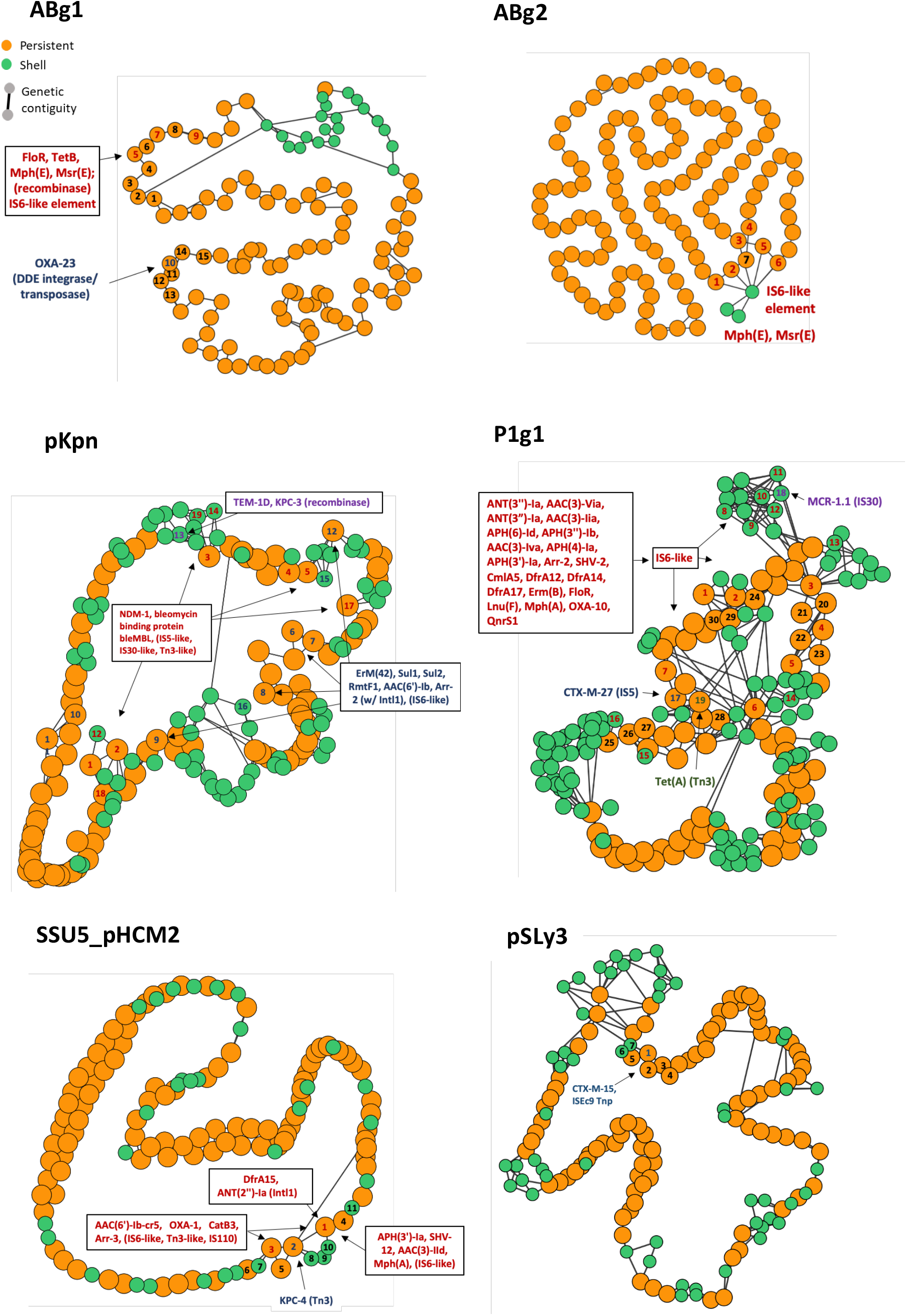
Genetic environments of ARGs in P-Ps’ pangenome graphs. Nodes are gene families. Genes (including ARGs) were groups into gene families (default parameters of PPanGGolin (43)) if they had a similarity of >80% identity covering at least 80% of the sequence. Orange = persistent, green = shell. Edges are shown for adjacent genes within the gene families (genetic contiguity). Gene families with colored numbers (red, blue, violet, green) are direct neighbors of ARG containing genetic elements (transposon, IS, integron, recombinase (separated by colors)). Black numbers are given for proximal gene families. **ABg1:** 1-3,6,8-9:hypothetical, 4:nucleoside, 5:ATPase AAA, 7:3’-5’ exonuclease pyrophospho-hydrolase. **ABg2:** 1,3:hypothetical, 2,7:ribonucleoside-diphosphate red. 4:toprim domain protein, 5:ATPase AAA, 7:3’-5’ exonuclease pyrophosphohydrolase. **SSU5_pHCM2**: 1:PhoH, 2-5, 7-11:hypothetical, 6:DNA ligase. **P1**: 1:SSB, 2-5,15,17,19-20,23-24,28: hypothetical, 6:cell division inhibitor (Icd-like), 7-11:tail fiber, 12:recombinase, 13-14:tail fiber assembly, 16:ResMod subM, 18:Ref family, 21:bleomycin hydrolase, 22:transglycosylase, 25:DNA repair, 26:Phd/ YefM (T-A), 27:doc (T-A), 29:lysozyme, 30:head processing. **pKpn**: 1:transcriptional regulator, 2,7,9,12-15,17-19:hypothetical, 3:phoH, 4:porphyrin biosynthesis protein, 5-6: AAA family ATPase, 8: ribonucl.-diphosphate reductase subunit 10-11:tail fiber domain-containing protein, 16:HsdR. **pSLyr3:** 1:DUF3927 family, 2:tellurite/colicin resistance, 3-7:hypothetical

Interestingly, we found that ARGs tend to be present in a small number of loci in the genomes of P-Ps. Within the P1-like and the pKpn-like genomes, the IS*6*-like transposable elements are inserted into a few distinct positions, whereas in genomes of SSU5-like and AB-like P-Ps all the insertions are concentrated in just one locus (figure 2). This is in line with our previous finding that P1 and pKpn genomes are more plastic than the average P-Ps (more complex, larger shell and cloud genomes, high number of plastic regions (23)). The conserved genes flanking the regions with ARGs often encode regulators, enzymes associated with DNA repair, or unknown functions. Few of them flank key highly conserved phage functions.

Overall, this analysis revealed diverse classes of ARGs in different groups of P-Ps (figure 2, supplementary figure S3-S7) that seem to have been acquired by the action of transposable elements.

### Integrons carrying ARGs are frequent in phage-plasmids

Class 1 integrons are not mobile by themselves, but plasmids often carry such integrons with ARGs (resistance integrons) (56). A recent analysis identified more than 1400 complete integrons in plasmids on the genome dataset used in our study (42). In contrast, no integron carried by a phage was reported so far. Accordingly, we searched for integrons in 3585 phages lacking evidence of being P-Ps and found no single integron in these elements. Since P-Ps have characteristics intermediate between plasmids and phages, we screened them for integrons. We found 27 integrons in P-Ps. Integrons were especially abundant in P1-like elements (n=11) (figure 3). Although, the SSU5 supergroup has the most members (n=268), just four P-Ps were predicted to have integrons in this set. Just in one P-P (NZ_CP016381), isolated from an *A. hydrophila* strain, two dissimilar integrons were detected. Furthermore, the *A. hydrophila* P-P has some similarity to P1 (wGRR = 0.07, 19 homologous genes), but not enough to class it as P1-like. Nine P-Ps with integrons were found in VP882-like and F116-like P-Ps.

**Figure 3:**
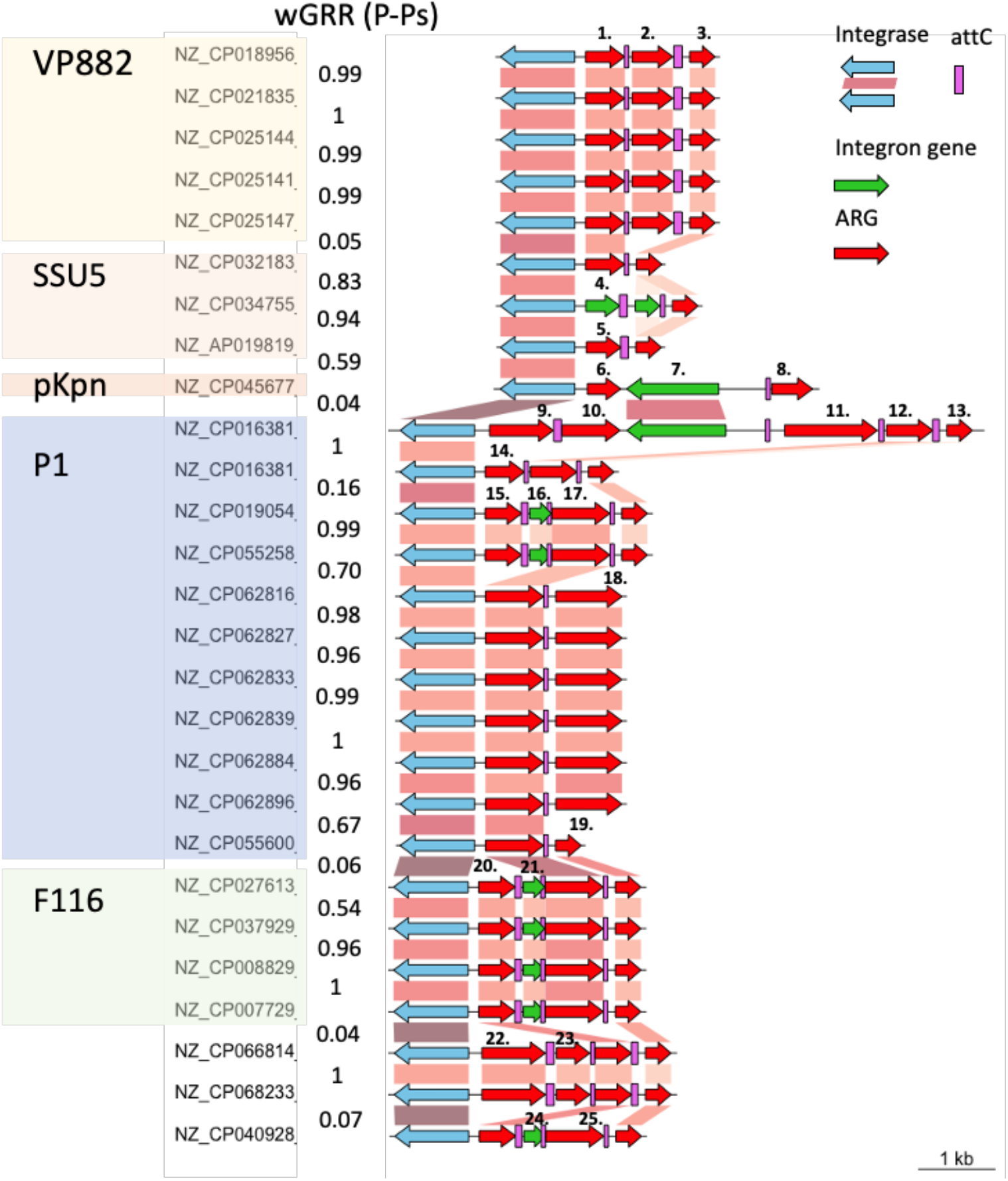
Integrons encoded in P-Ps. A. Genomic organization of integrons found in P-Ps, arranged by P-P groups and co-aligned by the class I Intl1 integrase. The P-P type is highlighted by differently colors. Gene-to-gene assignment is based on a blastp comparison when the alignment is at least 90% identical and covers at least 90% of the sequences. Blue gene arrows indicate the integrase gene, red arrows show AMR genes (99% identity, 99% coverage to reference sequences), and green arrows represent the rest of the genes within the integrons. Numbers indicated above non-homologous cassettes represent different types of ARGs: [1,14]:*ant(2’’)-Ia*, [2]:*aac(6’)-33*, [3,13,19]:*qacEΔ1*, [4]:*aac(3)-Ib*, [5]:*dfrA15*, [6]:*arr-2*, [7]:group II intron reverse transcriptase/maturase, [8]:*aac(6’)-Ib*, [9]:*bla*_CARB-2_, [10,17,25]:*aadA2*, [11]:*cmlA6*, [12]:*catB11*, [15,20]:*dfrA12*, [16,21,24]:DUF1010 protein, [18]:*aac(3)-VIa*, [22]:*bla*_GES-1_, [23]:*arr-6*.

These integrons have between two and five cassettes. Remarkably, nearly all genes within the cassettes were predicted to be ARGs (figure 3). As usual, *qacEdelta1* conferring resistance to antiseptics, was detected in most integrons (20/27) being part of the 3’ conserved segment. We found 15 co-occurrences of this cassette with that of *aadA2* (aminoglycoside nucleotidyltransferase) and 10 with those with *dfrA12*/*dfrA15* genes (trimethoprim resistance). A large diversity of other resistance genes was identified in integrons including aminoglycosides modifying enzymes (*ant(2’’)-Ia, aac(6’)-33, aac(3)-Via*), rifampicin resistance gene (*arr-2, arr-6*), chloramphenicol resistance (*cmlA6, catB11*) and different β-lactamases including the minor ESBL *bla*_GES-1_ and the penicillinase *bla*_CARB-2_ (figure 3). Hence, integrons in P-Ps encode a diverse panel of ARGs.

We compared the gene repertoire relatedness (wGRR) between integron-encoding P-Ps and the similarity between the integrons themselves. The gene cassette arrays tend to be very similar when they are in the same type of P-P and very distinct across unrelated P-Ps. However, in a few cases (pointed out by black arrows in figure 3), dissimilar P-Ps have similar integron cassettes (>90% identity and 90% coverage) suggesting an epidemic spread of genes providing a selective advantage (resistance to aminoglycosides and antiseptics). In all cases, the integrons of P-Ps had very similar IntI1 type integrases. These results suggest that, like it is the case for plasmids, type I integrons act as reservoirs for multiple ARGs in P-Ps.

### Phage-plasmids with resistance genes are induced by mitomycin C

The recent acquisition of ARGs in P-Ps might make them inactive phages, as previously observed in a plasmid resembling a P1-like element (25), either because the insertion inactivates relevant functions or because natural selection of the bacterium could select for inactive elements. To test if some ARG-encoding P-Ps are functional phages, we screened a collection of draft genomes of CRE strains for ARG-encoding P-Ps. We identified six strains (two *E. coli* and four *E. cloacae*) that we sequenced using long reads (supplementary figure S8). These genomes had six P-Ps with ARGs: one P1-like (of 163A9) in *E. coli*, and five SSU5-like P-Ps, one from *E. coli* (of 166F4) and four from *E. cloacae* (169C2, 170E2, 171A5, 211G7) (supplementary table S2). We then tested if these P-Ps are induced by DNA damage by exposing the bacterial cells to mitomycin C (MMC). 3-4h after MMC addition, a drop of cell-density indicated the phage-dependent cell lysis (caused by SOS response and consecutive prophage and/or P-P induction) (supplementary figure S9). Phage particles were purified, their DNA extracted and sequenced by short reads (after digestion of chromosomal DNA (gDNA), see methods). We then conducted hybrid assemblies by combining the short reads from the MMC experiment with the long reads from the genome sequencing (figure 4A).

**Figure 4:**
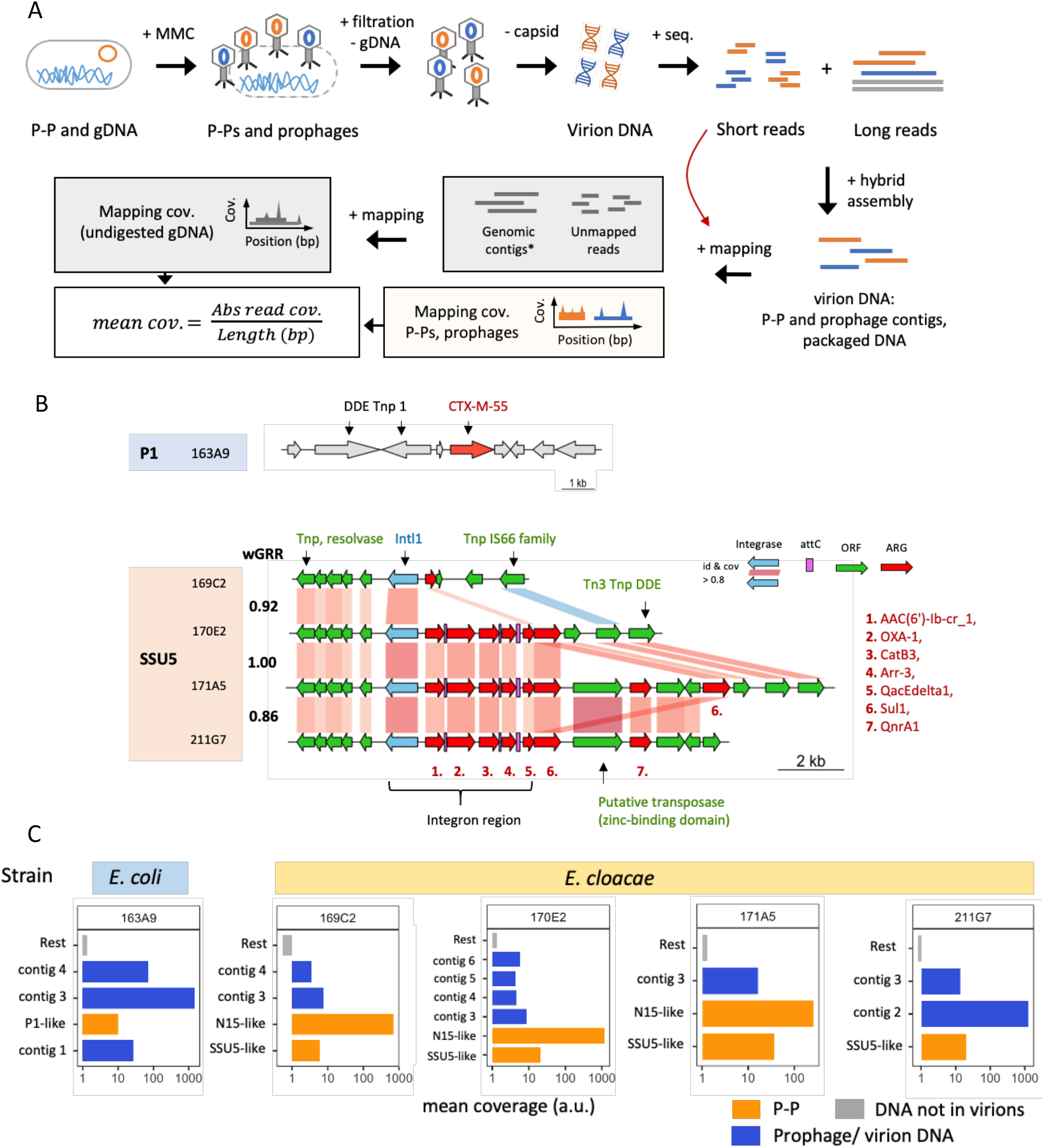
Induction of P-Ps and prophages in CRE strains. **A**. CRE strains w/ ARG-encoding P-Ps were induced by 5 μg/ml MMC. 4h after induction, phage particles were purified and chromosomal DNA (=gDNA) removed by DNase I digestion. The phage capsid was degraded by proteinase K, virion DNA was purified and sequenced. The obtained short reads were co-assembled with long-reads from the genomic sequencing experiment (see methods). The assemblies were compared to P-P and phage genomes, subsequently assigned and the read mapping coverage was computed (by mapping the short reads from the MMC experiment on them). The reads that did not map the assemblies were used to compute the background coverage caused by undigested gDNA (by mapping on genomic contigs obtained by the long-read assembly). **B**. In the genome of the P1-like P-P of the *E. coli* 163A9 the CTX-M-55 gene is found next to two DDE transposases. The ARGs encoded in the SSU5-like P-Ps from the *E. cloacae* strains 169C2, 170E2, 171A5 and 211G7 are in a complex region containing transposases and integrons. Homology assignments between P-Ps were done when sequence similarity was of at least 80% identity and covered 80% of the sequence of the gene (retrieved from an all-vs-all blastp comparison). The similarity between P-Ps is indicated by the weighted gene repertoire relatedness (wGRR). **C**. Average read coverage was obtained and calculated as described in A. Shown are all contigs larger than 10kb. The coverage (a.u. = arbitrary unit) is plotted on a logarithmic x axis for P-P contigs (orange), contigs assigned to prophages or virion loaded DNA (blue) and average background coverage (rest coverage) obtained after mapping the remaining reads on genomic contigs (grey) for each tested CRE strain.

For all P-Ps, except of strain 166F4, the co-assembly resulted in closed circular contigs or in larger assemblies with good homology to known P-Ps (supplementary figure S10-S11), that confirmed the structure of the replicons. This opened the possibility of studying the genetic context of the ARGs in these P-Ps. The P1-like P-P of the *E. coli* strain 163A9 contains two DDE transposases next to the β-lactamase encoded by a *bla*_CTX-M-55_ gene (figure 4B). The four SSU5-like P-Ps from *E. cloacae* are very similar (wGRRs from 0.86 to 1, figure 4B, supplementary figure S11) and their ARGs are in the same loci, in a complex region including various transposases and a type 1 integron with a very similar integrase (identity and coverage >80%) (figure 4B). Three P-Ps contain integron regions with five ARGs, whereas the one from the strain 169C2 has a very similar integrase gene but lacks gene cassettes (figure 4B). In addition, downstream of the integron region a few (one to three) more ARGs are in a locus with multiple transposases. The number of ARGs of the SSU5-like P-Ps ranges from one (169C2) to eight (171A5) and they are predicted to encode resistance against several antibiotics including penicillins (*bla*_OXA-1_), fluoroquinolones (*qnrA1*), aminoglycosides (*aac(6’)-Ib*) or sulfonamides (*sul1*) (figure 4B, supplementary table S2).

To verify that P-P DNA is found in the virions, we mapped the short reads from the MMC experiment on the co-assemblies and retrieved the average mapping coverage (figure 4A). Unmapped reads were extracted and used to obtain the relative frequency of non-digested chromosomal DNA (=background signal) by mapping them on the genome contigs (see methods) (figure 4A). We considered that highly covered P-Ps and prophages, relative to the background chromosome, were induced and packaged into virions. Five of the six tested strains were indeed induced and produced viral particles with the ARG-encoding P-Ps (figure 4C). The DNA of these elements was present at diverse frequencies, possibly a result of different burst sizes (which might be further affected by the presence of other prophages (57)). For one of the six strains (*E. coli* 166F4), we did not obtain any reads mapping the P-P contig suggesting that this SSU5-like P-P is inactive or not inducible by MMC under our experimental conditions. Interestingly, in the genomes of three strains (*E. cloacae* 169C2, 170E2 and 171A5) we found two different types of P-Ps: the SSU5-like encoding the ARG and another P-P lacking ARGs that is related to the N15 group. Noteworthy, we could also assign some of the sequences retrieved from the viral particles to chromosomal loci with prophages in the CRE strains, which shows that they were also induced by MMC (figure 4C, supplementary table S3). The coverage of P-Ps and integrative prophages in the analysis of the viral particles is typically at least one order of magnitude higher than the average coverage of the background, showing that the result is not due to random contamination by bacterial DNA. Hence, most of the analysed P-Ps, with or without ARGs, are inducible and are packaged into virions.

The MMC induction experiments confirmed that P-Ps with ARGs are inducible and packaged into virions. Hence, these P-Ps confer resistance to antibiotics to the bacterium and function as real phages. Here, we test whether they are also capable of infecting, lysogenizing, and converting other host strains into becoming antibiotic resistant. Phages tend to have narrower host ranges than conjugative plasmids, possibly because of their requirement for specific receptors at the cell envelope (58), the existence of numerous bacterial defenses against phages (59), and the presence of other prophages (60). This results in phage-bacteria interaction matrices that tend to reveal a very low frequency of productive infections for temperate phages (61, 62). Hence, the first challenge was to identify permissive hosts different from the strain carrying the P-P.

The four SSU5-like PPs are very similar and presumably have similar host ranges. Since the requirements of host range of these phages in this host are poorly understood, we tested 18 different *E. cloacae* strains retrieved from the Pasteur and the German collections (DSMZ) (supplementary table S4). However, no lysogenic conversion was observed by the SSU5-like P-P.

We chose a diverse panel of 12 *E. coli* and one *S*. enterica (CIP 82.29T) strains from the Pasteur collection to infect with the P1-like P-P (supplementary table S4). The *S. enterica* strain was of particular interest to test the host range of the P-P. We then conducted infection experiments, where we purified the phage particles, incubated them with the potential host strains and screened for resistant lysogens by plating the mixture on antibiotic-containing plates (here carbenicillin, see Methods). For the *S. enterica* strain we did not obtain lysogens. However, we found four different *E. coli* recipient strains (55989, CIP 105917, CIP 53.126, CIP 76.24), all from phylogroup B, that were initially sensitive but became resistant to carbenicillin after the infection with the P1-like P-P of strain 163A9 (figure 5AB). The sequencing of the genomes of the recipient strains confirmed the acquisition of the ARG and the rest of the P-P (figure 5CDE, supplementary figure S12). Moreover, susceptibility tests with various β-lactam antibiotics (broad-spectrum penicillins, cephalosporins of different generations, carbapenems) confirmed the presence of the CTX-M-55 ESBL in the lysogenized strains. All four strains show resistance to three broad spectrum penicillins (Ticarcillin, Piperacillin, Amoxicillin) and a 3^rd^ generation cephalosporin (Cefotaxime) (supplementary figure S13). Finally, we tested if the P-P is fully functional, by testing if it can be induced in the new host and used to infect another cell. We exposed the *E. coli* 55989 strain with the P1-like P-P to MMC, purified the particles and used them to infect the original antibiotic-sensitive *E. coli* 55989 strain. This revealed the acquisition of the P-P and the lysogenic conversion (supplementary figure S14), thereby confirming that it is a fully functional phage. This demonstrates that natural P1-like P-Ps can transfer ARGs as phages and result in lysogenic conversion of other strains.

**Figure 5:**
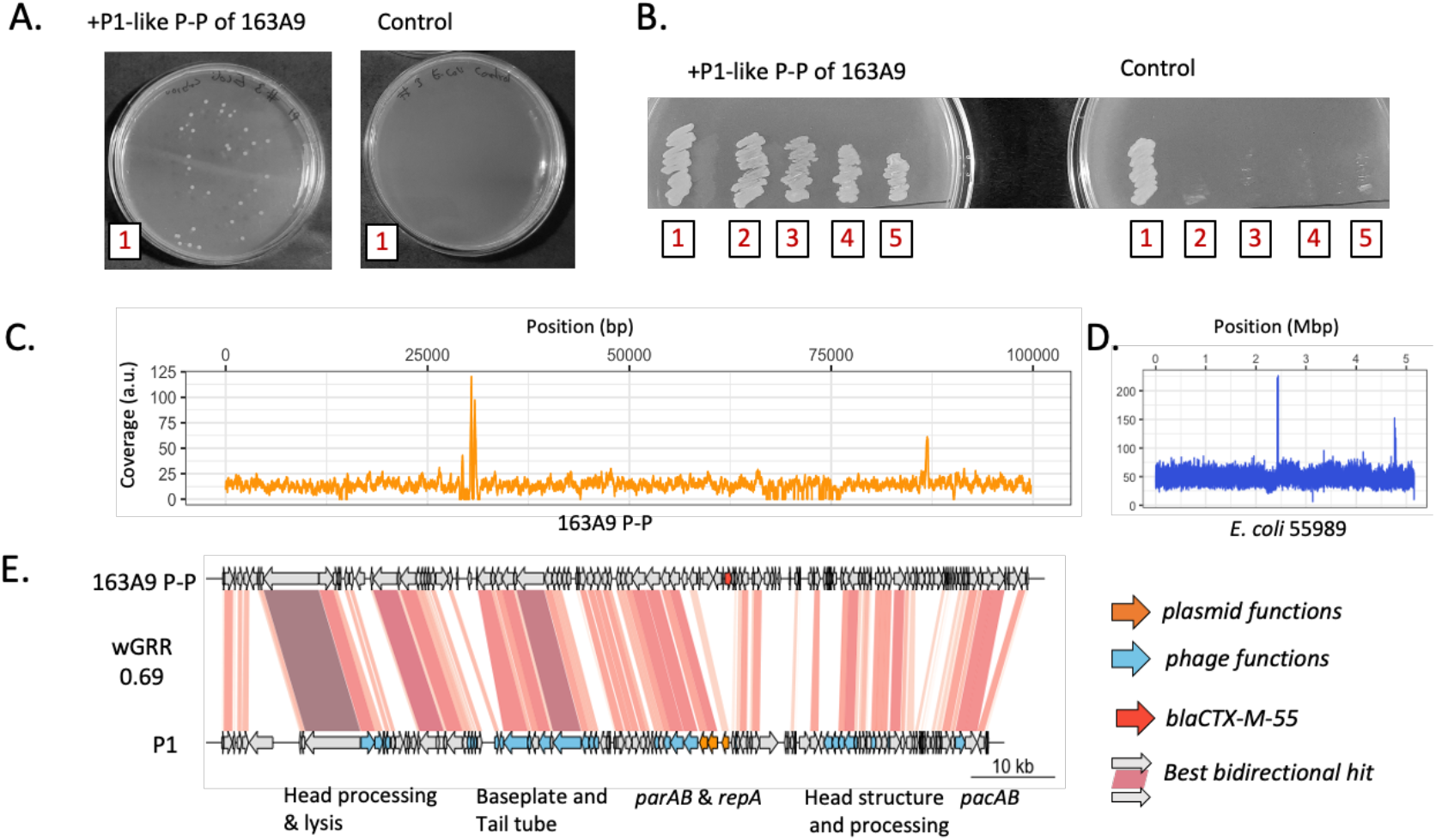
Lysogenic conversion of different *E. coli* (55989 [2], CIP 105917 [3], CIP 53.126 [4], CIP 76.24 [5]) by the P1-like P-P of strain 163A9 [1]. **A + B**. After the infection and plating experiment four tested *E. coli* strains acquired resistance to carbenicillin. Examples of colonies of strain 55989 with the P1-like P-P of 163A9 are shown on LB agar plates with carbenicillin 100 ug/ml (A). The original host of the P1-like P-P and all four lysogens are resistant to ampicillin 100 ug/ml (B). **C & D**. Their genomes were isolated and sequenced by short reads (supplementary figure 12). Shown is the read coverage for the P-P genome (C) and the chromosomes of the host strain *E. coli* 55989 (D). **E**. Genome comparison of the P-P from 163A9 and P1 and are co-aligned to the first gene of P1. The alignment is matched with the read coverage plot in C. The function of P1 genes were retrieved from Łobocka et al (63).

## Discussion

ARGs and integrons are commonly found in plasmids (8), but very rarely in phages (16). P-Ps are temperate phages and plasmids. Here, we show they are more much likely to encode ARGs than the other phages. We also show that they frequently encode integrons with ARGs, which had never been observed in functional phages. We demonstrated that one of the P-P was a fully functional phage that could be induced, produced lysogens resistance to broad-spectrum cephalosporins, and this could be shown for two full cycles of induction, infection and conversion. Nevertheless, there are some limitations to our study. Some P-Ps may be inactive, a trait common among integrative prophages (64). Alternatively, some P-Ps may not be inducible by MMC. Still, we tested six strains that had nine putative P-Ps and all but one could be induced to produce viral particles containing phage DNA. This suggests that many P-Ps are still functional. We could not obtain new lysogens for a group of closely related SSU5-like P-Ps by screening for antibiotic resistance. To identify ARGs, we applied very strict criteria (99% identity over 99% sequence coverage), however their expression may depend on the genetic background. The lack of lysogens may also be caused by bacterial resistance to phage infection. Previous works have shown that interaction matrices of bacteria with temperate phages tend to be very sparse (61), i.e. most combinations do not result in productive infection. This is because many bacteria have anti-phage systems, lack appropriate cell receptors, or have phages repressing infecting P-Ps (60). In addition, the replication initiators of P-Ps may be incompatible with those of resident plasmids, preventing their establishment in lysogens. Further work will be needed to explore the host range of P-Ps carrying ARGs. In contrast, we found multiple *E. coli* strains susceptible to the P1-like phage of strain 163A9. These differences may be associated with P-Ps host range, which is known to be unusually broad in P1 (65, 66).

A few previous reports identified ARGs in P-P-like elements among enterobacteria, even though evidence of induction is often lacking (25, 26, 67). In our study, we show that this is a general trend of P1-like elements but also of different other types of P-Ps. Such cases can be found in other bacterial clades of important nosocomial pathogens, e.g., in *Acinetobacter*. Overall, they are much more likely to carry ARGs than the other phages. We show that they carry a wide diversity of resistances. Most worrisome, many clinically relevant ARGs are found in P-Ps, including the carbapenemase genes *bla*_KPC-3_ and *bla*_NDM-1_. The *bla*_KPC_-like genes are involved in the diffusion of carbapenem resistance in Italy, Israel and USA, whereas the *bla*_NDM_-like gene is disseminated worldwide (68). Among the last-resort antibiotic, colistin was reintroduced in the armamentarium to fight against carbapenem-resistant Gram-negative rods despite its nephrotoxicity. Plasmid-mediated colistin resistance has been recently described (69). This ARG, named *mcr-1*, was also identified in P-Ps. Beyond these two important resistance mechanisms, the *rmtF* gene is of further importance and also carried by P-Ps. This gene encodes a 16S RNA methyltransferase conferring resistance to almost all aminoglycosides used for treatment (70). Thus, P-Ps are involved in the diffusion of resistance to the main antibiotic families including β-lactams (e.g. *bla*_KPC-3_), aminoglycosides, fluoroquinolones (e.g. *qnrA1*) and polymyxins.

The presence of ARGs genes in P-Ps raises concerns that are common to the resistances found in other plasmids, notably that they can spread these genes across bacterial populations. However, the fact that P-Ps are also phages raises additional concerns. First, and unlike conjugative plasmids, phages transfer their DNA in viral particles and do not require direct contact between cells for the transfer. Hence, they can transfer between bacteria present in different time and places. Second, the lytic cycle of phages amplifies their genomes hundreds of times (e.g. 400 for P1 (71)) for packaging in the viral particles, which may result in bursts of transfer of ARGs. The process of phage replication in the cell could also lead to over-expression of ARGs and liberate enzymes that detoxify the environment for the remaining bacteria, a process akin to the production of the Shiga toxin from the prophages encoding it (72). However, it must be stated that in our experiences the only drug that induced P-Ps was MMC. Finally, P-Ps are more likely to recombine with other phages than the remaining plasmids, because they share numerous homologous genes (23). This may pose a threat of ARG transfers to other phages.

Plasmids are known to contain many transposable elements and integrons that facilitate the translocation of ARGs within and between replicons (9). In contrast, phages typically have very few if any such elements (73). Here, we show that P-Ps have numerous transposable elements associated with ARGs and integrons. Hence, P-Ps can take advantage of genetic elements typical of plasmids, transposable elements and integrons, to acquire ARGs that can then be spread horizontally by viral particles. Integrons are reservoirs of ARGs and, especially in clinical settings, promote the spread of multi drug resistances (74). Their identification in P-Ps is worrisome, because the integron ability to incorporate novel cassettes from other integrons implies that upon acquisition of an integron the repertoire of ARGs of the P-Ps can evolve faster to incorporate novel types of resistance.

This raises the question of why P-Ps have so many more ARGs than the other phages. About half of the sequenced phages are virulent (75). They are not expected to carry ARGs because they do not produce lysogens, although this cannot be completely excluded, since they may produce pseudo-lysogens (76). The causes for the different frequency of ARGs in P-Ps and other temperate phages are less obvious. It could be argued that the frequency of ARGs in P-Ps was caused by their abundance in bacterial pathogens. However, most sequenced phages in the database are also from a few genuses including many enterobacteria, and this does not seem sufficient to explain the difference in the number of ARGs present in the two types of elements. For example, the naturally-occurring phages of *E. coli* used this study, which were not P-Ps, are completely devoid of identifiable ARGs. We propose that differences between P-Ps and integrative, temperate phages result from a combination of factors. First, P-Ps tend to be larger than the other phages (23). This is particularly relevant for phages, because they can only package an excess of a few percent of their genome size. A sudden larger increase in genome size precludes packaging of the genome and thus blocks phage transfer. Hence, a genome of 49 kb, like phage lambda, can only accommodate very small insertions. In contrast, P1-like elements are on average 96 kb (23), and can tolerate larger changes. Since the integration of ARGs in P-Ps involves the transposition of the gene and flanking transposable elements and/or integrons, these insertions may be too large for most integrative phages. Accordingly, we found ARGs in the largest P-Ps, like P1-like and SSU5-like, but not in the much smaller N15-like elements (average 55 kb, (23)). Differences between P-Ps and other phages could also be caused by the regions of homology to plasmids of the former. This might facilitate genetic exchanges between plasmids and P-Ps.

Independently of the reasons leading to an over-representation of ARGs in P-Ps relative to the other phages, the subsequent evolution of the loci containing them in P-Ps may result in streamlined compact loci that could be easier to transfer to other phages. Indeed, integrative temperate phages already encode many virulence factors, and we cannot find a reason why, given enough time, they will not eventually acquire ARGs. This would be a most worrisome outcome of the recent evolutionary process of acquisition of ARGs by human-associated bacteria, since, as stated above, phages are extremely abundant, spread very fast in the environment, and can infect bacteria in different geographical locations and time.

## Supporting information

supplementary_tbl_s1

supplementary_tbl_s2

supplementary_tbl_s3

supplementary_tbl_s4

supplementary_tbl_s5

## Acknowledgements

INCEPTION project (PIA/ANR-16-CONV-0005), Equipe FRM (Fondation pour la Recherche Médicale): EQU201903007835, Laboratoire d’Excellence IBEID Integrative Biology of Emerging Infectious Diseases [ANR-10-LABX-62-IBEID], SALMOPROPHAGE ANR-16-CE16-0029. Sequencing was done at the Biomics Platform, C2RT, Institut Pasteur, Paris, France, supported by France Génomique (ANR-10-INBS-09) and IBISA. We thank especially Marc Monot, Goerges Haustant, Laure Lemée, Valérie Briolat, Laurence Ma, Elodie Turc, Amandine Buffet and Matthieu Haudiquet for providing support, advice and services in DNA isolation, library preparation and sequencing. This work used the computational and storage services (TARS & MAESTRO cluster) provided by the IT department at Institut Pasteur, Paris.

## Table legends

Table legends

Supplementary table 1:

Antibiotic resistance genes detected in phage-plasmids.

Supplementary table 2:

Phage-plasmids with ARG in CRE strains.

Supplementary table 3:

VirSorter2 and wGRR analysis of co-assemblies retrieved after MMC-induction experiment (including average read coverage).

Supplementary table 4:

Bacterial strains that were used for infection experiments with ARG encoding P-Ps.

Supplementary table 5:

Accession numbers on the reads and assemblies from the genome sequencing, MMC induction (hybrid assembly included) and re-sequencing of lysogens.

