## supplementary_file for "Phage-plasmids spread antibiotic resistance genes through infection and lysogenic conversion"

### Supplemental Figures

A.

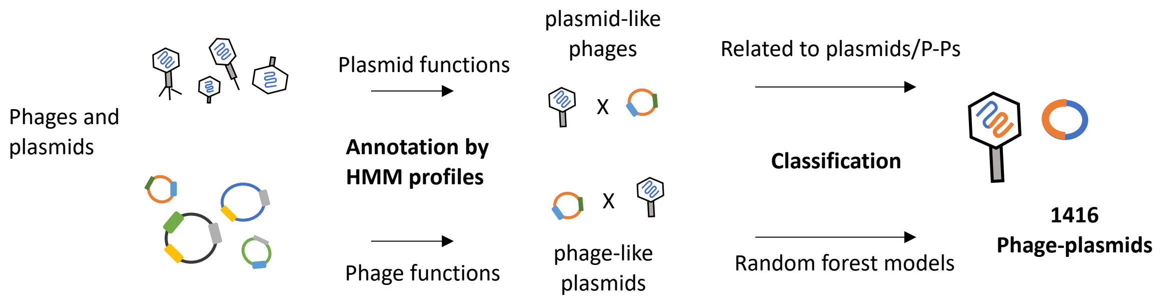

B.

| Database version | AB-group | SSU5-supergroup (pCAV, pKpn, pMT1, pSLy3, SSU5_pHMC2) | P1-group | N15-group |
|---|---|---|---|---|
| Mai 2019 | 24 | 145 (8+38+35+29+35) | 64 | 42 |
| March 2021 | 40 | 268 (14+94+43+62+55) | 149 | 71 |

Figure S1: Identification of phage-plasmids.
**A.** 25275 sequences of phages and plasmids, that were retrieved from RefSeq in March 2021, were searched for functions of P-Ps (as in a previous approach (1), see Methods). First, phage functions were annotated in plasmids and, secondly, the annotation profiles were evaluated by random forest classification models. In phages, we initially searched for plasmid features (partitioning, replication), and subsequently compared positive cases to known P-Ps and plasmids. All phages with plasmid genes having a higher similarity than wGRR = 0.4, were added to the P-P list. Overall, 1416 P-P were detected (including the P-Ps found in the first approach (1). **B.** All new detected P-P sequences were searched for homology to genomes of known P-Ps and subsequently assigned to corresponding P-P groups (AB, SSU5 supergroup and groups, P1 group, N15 group). Assignment was done, when a P-P had a wGRR $\geq$ 0.5, based on at least 50% of the genes, to a grouped P-P.

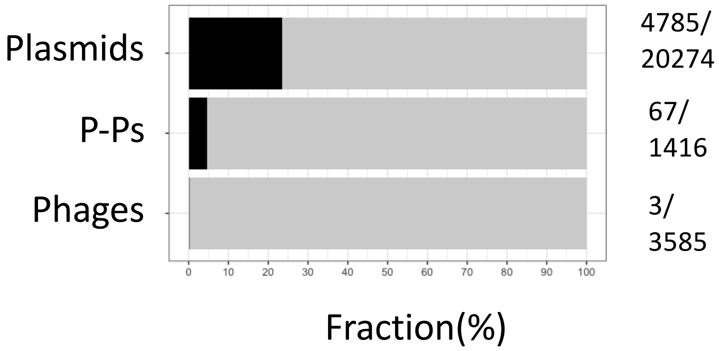

Figure S2: ARG-encoding plasmids, phage-plasmids and phages detected by AMRFinderPlus (2).

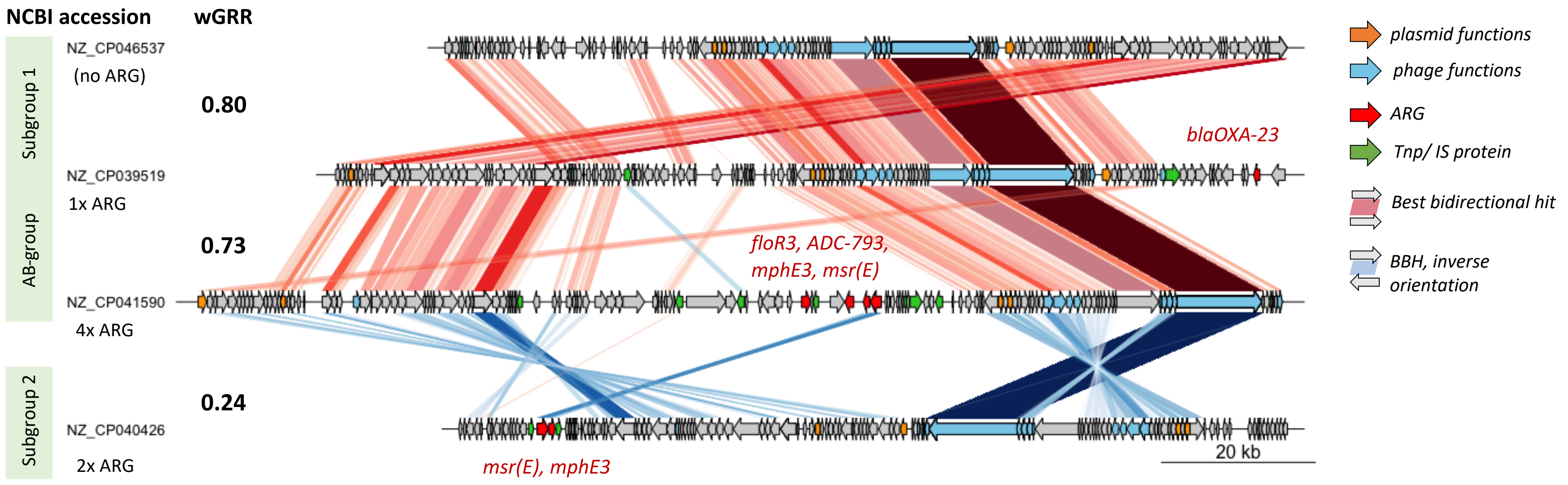

Figure S3: P-Ps of the AB subgroup1 and subgroup2 with and without ARGs.

Genome-to-genome comparisons of P-Ps with and w/o ARGs. wGRR relatedness is indicated between the genomes. Assigned regions (in red=directed, blue=inverse orientation) are showing best-bidirectional-hits (BBHs) that are based on an all-vs-all protein sequence comparison (by MMseqs2 (3)) with at least 35% identity, 50% sequence coverage and an e-value =<1e$^{-4}$. Plasmid genes (partition and replication functions) are orange, phage functions (structure, lysis, packaging) shown in blue, ARGs in red and genes assigned to transposases/IS elements in green arrows. Names of ARG are shown in the order they appear in red.

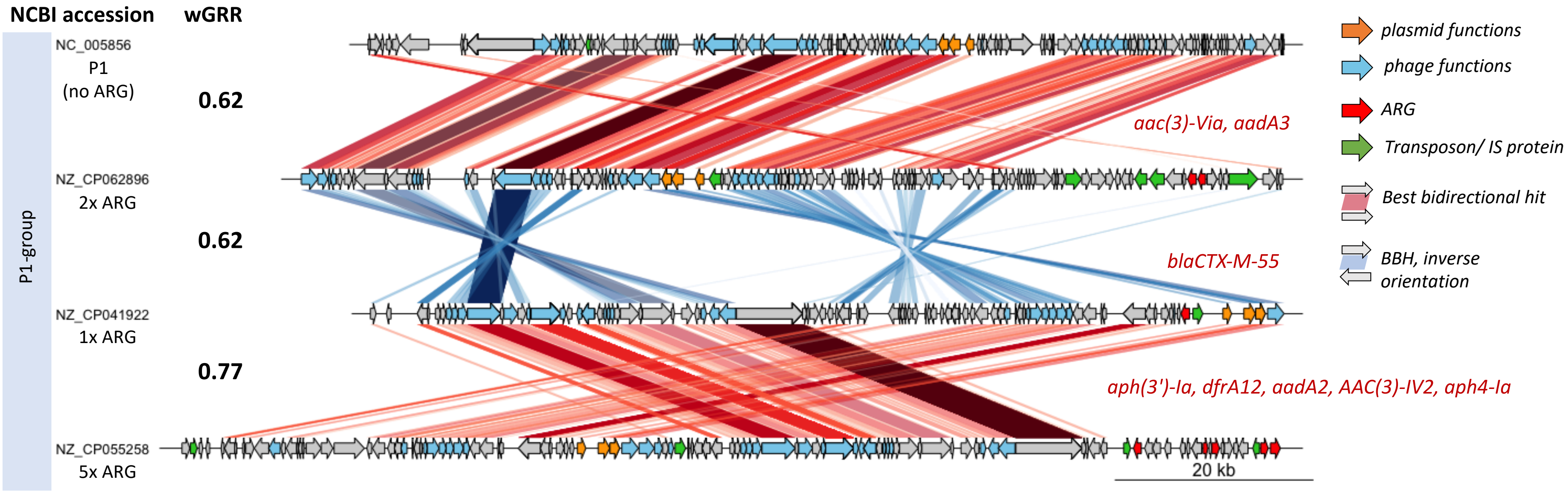

Figure S4: P-Ps of the P1 subgroup 1 w/ and w/o P-Ps.
Same as described in Fig S3.

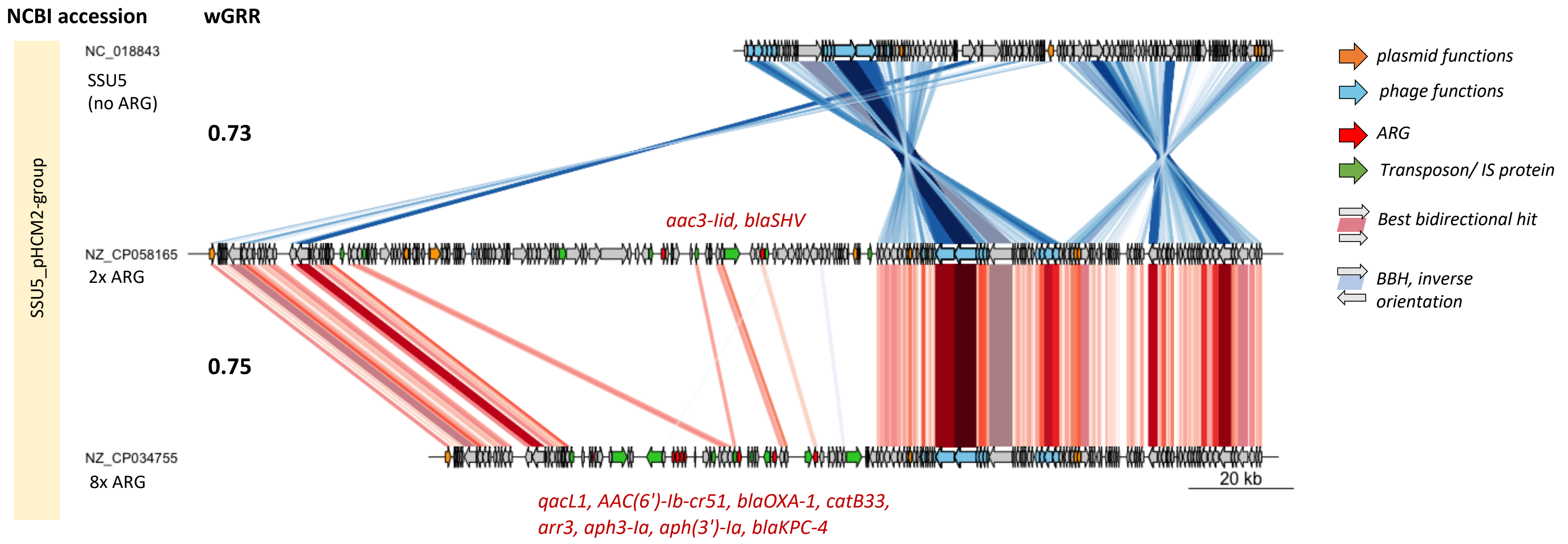

Figure S5: P-Ps of the SSU5_pHCM2 group with and without ARGs.
Same as described in Fig S3.

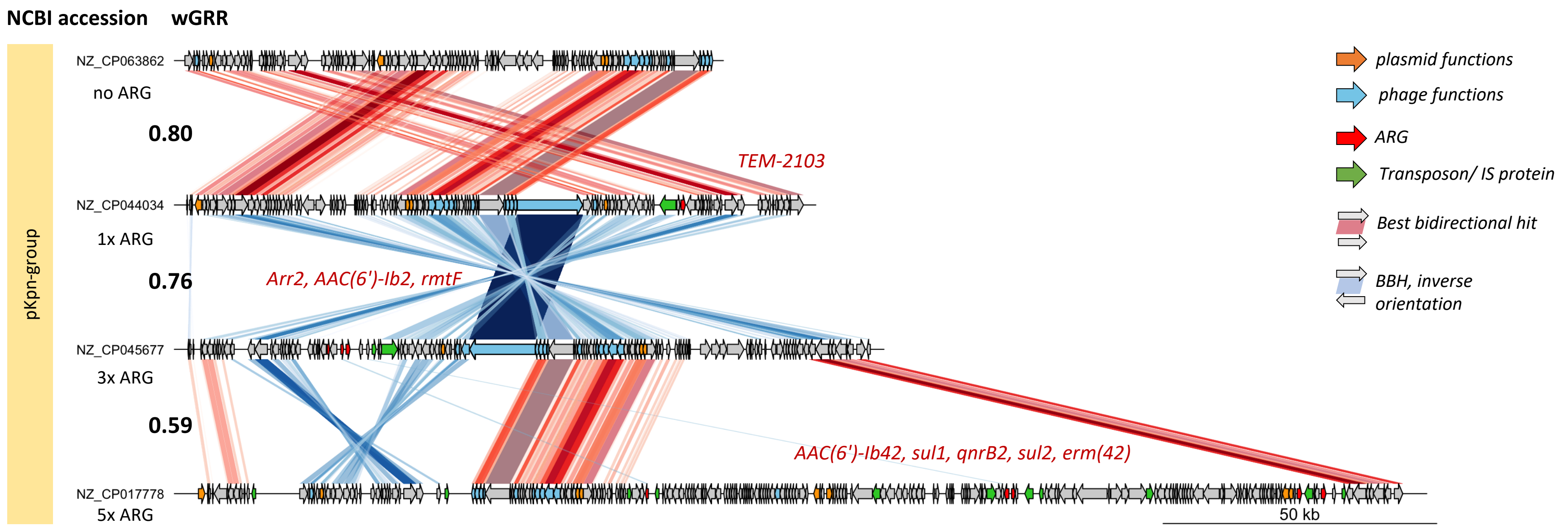

Figure S6: P-Ps of the pKpn group with and without ARGs.
Same as described in Fig S3.

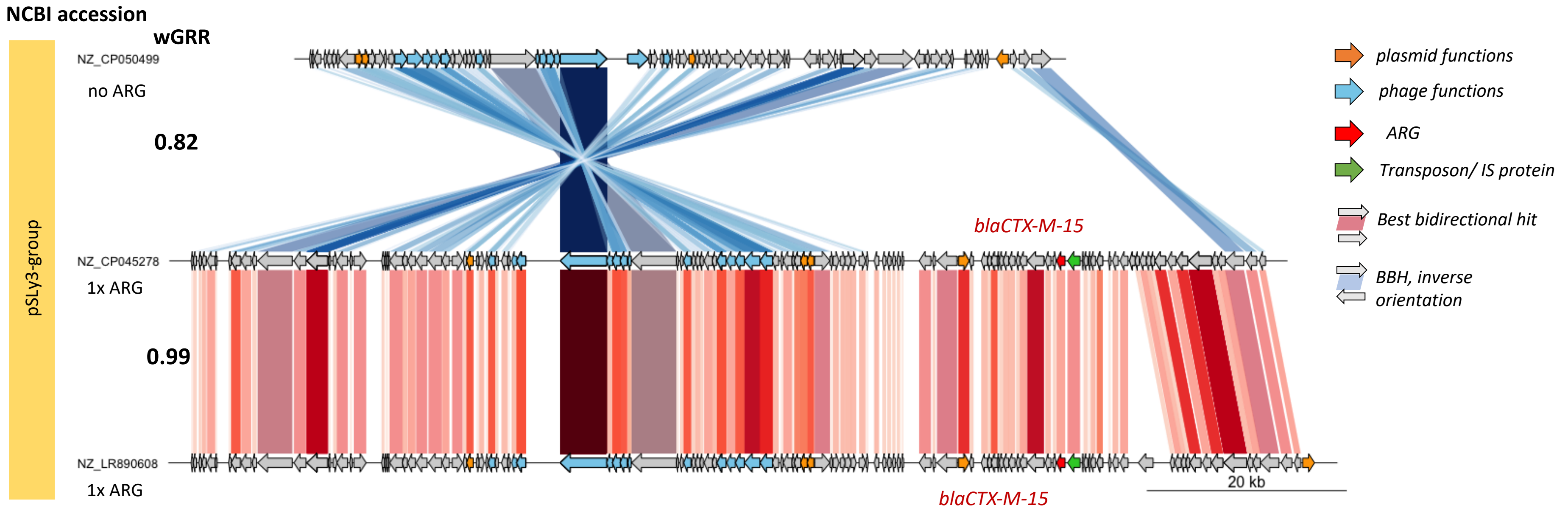

Figure S7: P-Ps of the pSLy3 group with and without ARGs.
Same as described in Fig S3.

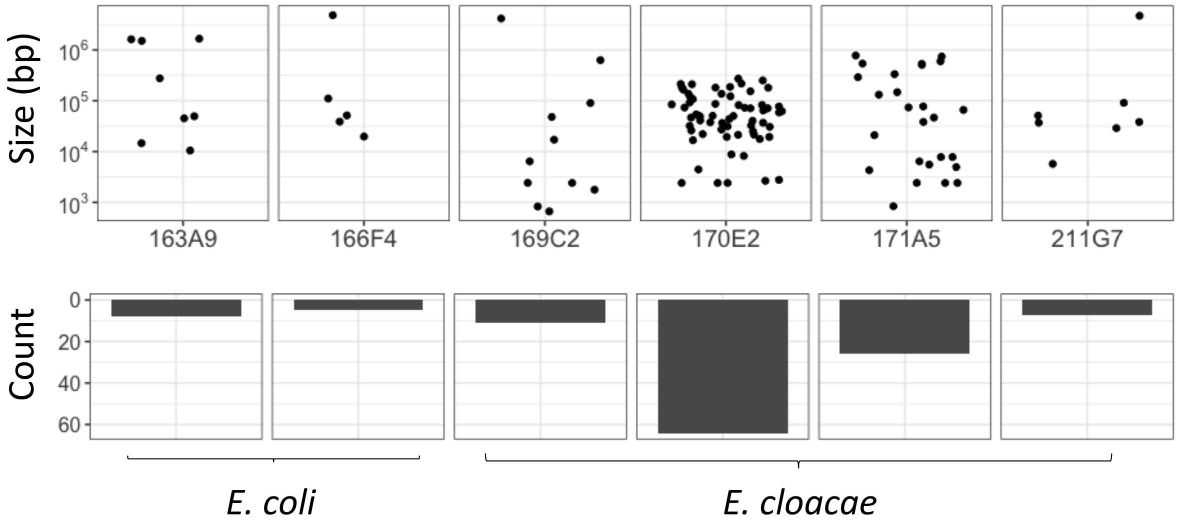

Figure S8: Number and length of assembled contigs of CRE strains with P-Ps.

The genomic DNA of the CRE strains with ARG-encoding P-Ps (163A9, 1664F4, 169C2, 170E2 171A5 and 211G7) was isolated and sequenced by the PacBio long-read technology. The obtained reads were assembled with flye (4) using default parameters. The scatterplots (upper panel) show the lengths of the contigs per strain and the overall number is represented in bar plots (lower panel).

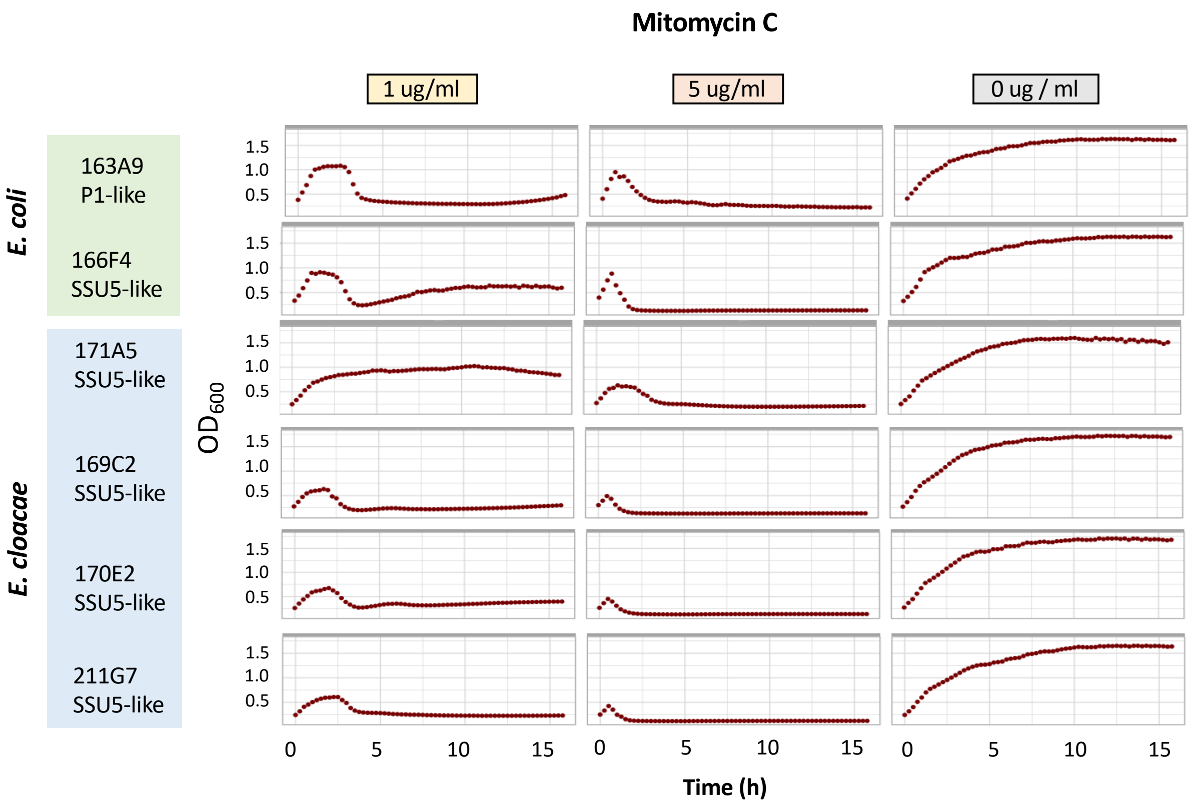

Figure S9: Growth experiments of P-P carrying CRE strains in LB with mitomycin C.
All CRE strains with P-Ps that have AMR genes were tested for inducibility by exposing them to 1 µg/ml and 5 µg/ml MMC in comparison to the control (no MMC). The growth experiments were conducted in 96-well plates and each strain was tested twice (technical replicate). After the addition of MMC, growth was followed by measuring optical densities at 600 nm by a plate reader under shaking conditions and 37 °C.

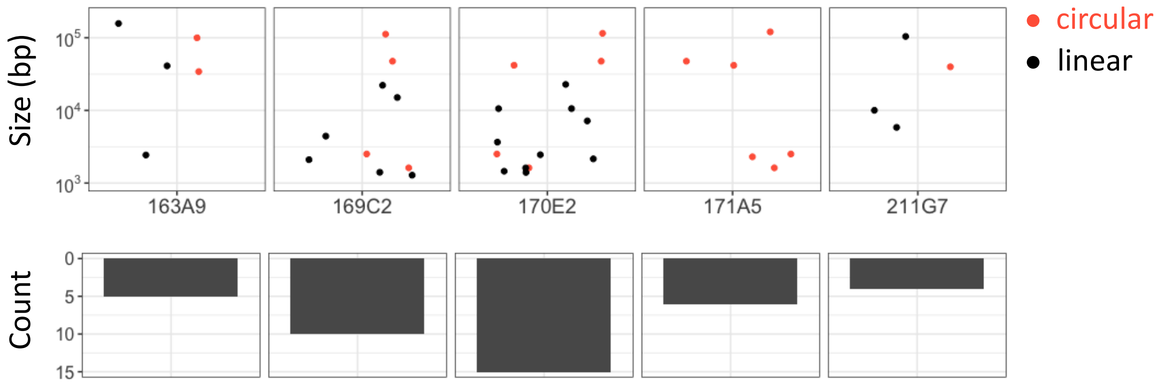

Figure S10: Number, length and topology of contigs assembled using an hybrid method after sequencing the DNA found in virions.

Short-reads of the MMC induction experiment and long-reads of the genome sequencing were used to conduct a co-assembly with Unicycler (5). Contig lengths are shown in the upper panel in the dot plots, circular contigs in red, linear or unassigned contigs in black. The total contig number per strain is represented in bar plots in the lower panel.

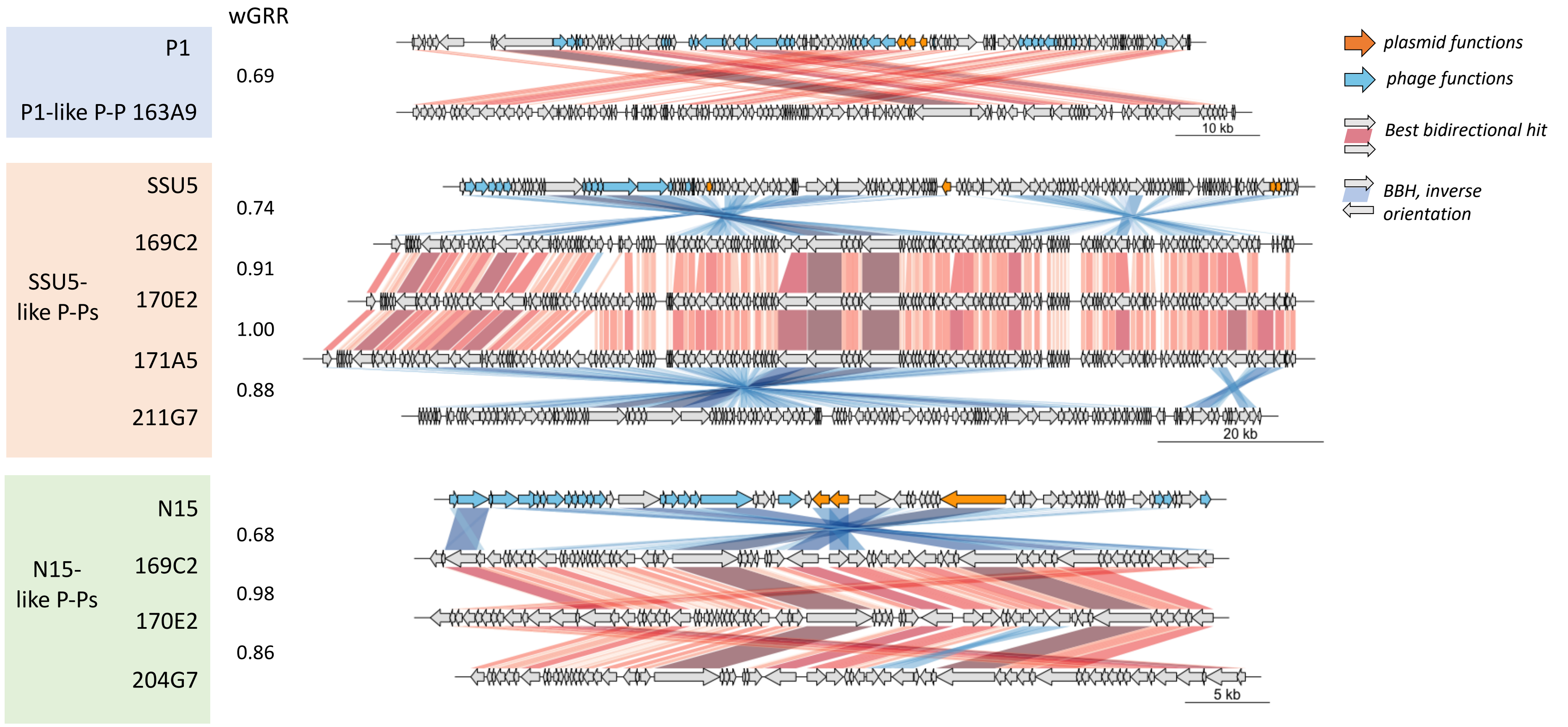

Figure S11: Comparisons of P-Ps from CRE strains with known P-Ps.

Plasmid functions (partitioning, replication) are shown in orange and phage genes (structure, packaging, lysis) in blue arrows. Assignment are BBHs that are based on an all-vs-all protein sequence comparison (by MMseqs2) having at least 35% sequence similarity, 50% coverage (both sides) and an e-value =< 0.001. BBHs in the same orientation are shown in red, inverse in blue assignments.

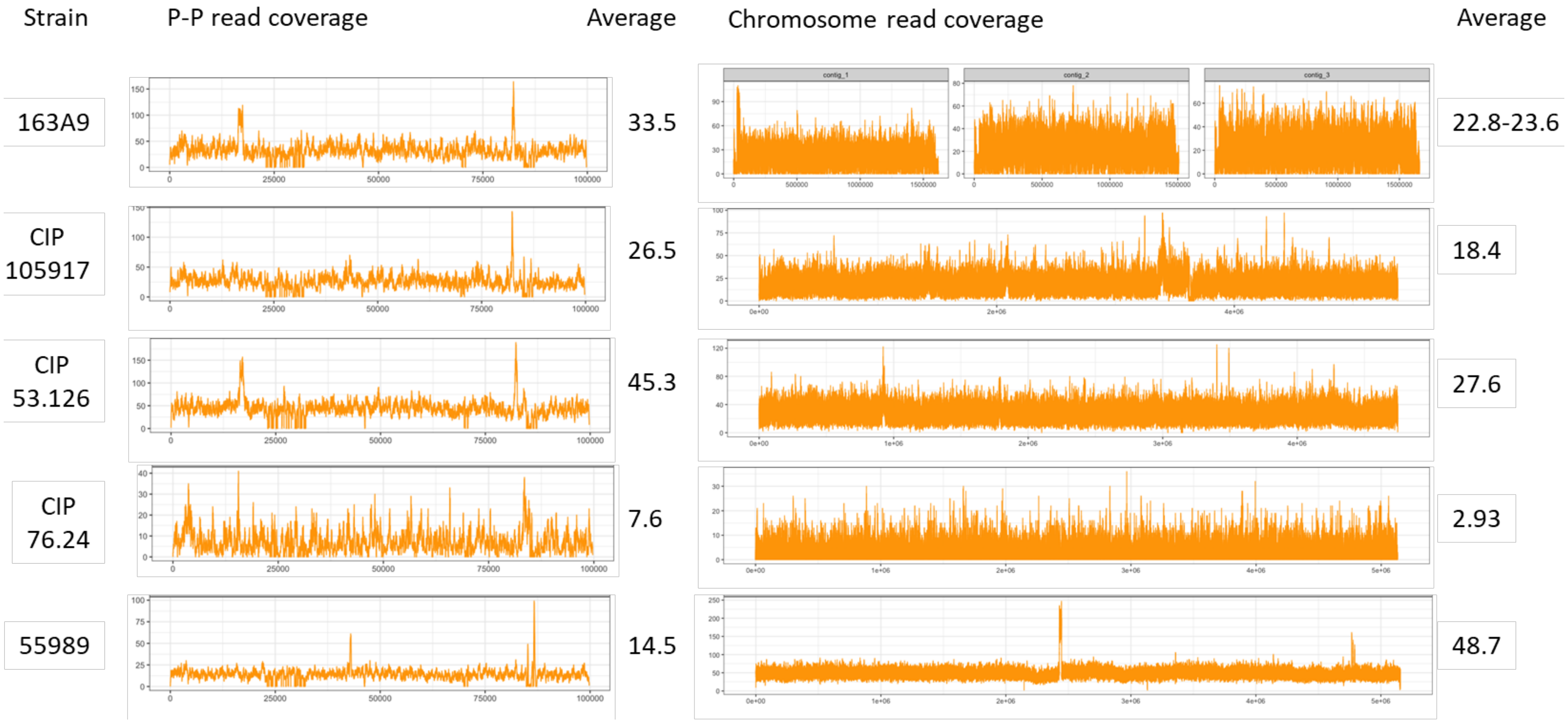

Figure S12: Genome re-sequencing of lysogens with P1-like P-P of strain 163A9.

Genomes of all *E. coli* lysogens (CIP 105197, CIP 53.126, CIP 76.24, 55989) having the P1-like P-P isolated from CRE strain 163A9 (including the original strain 163A9) were isolated and sequenced by short reads (Illumina). Shown is the read coverage for the P-P genome and the host chromosome (55989: NC_011748, CIP 105917: NZ_CP041623, CIP 53.126: NZ_CP022959, CIP 76.24: NZ_CP009072). For the CRE strain 163A9 the mapping was done on the three largest assembled contigs (contig 1: 1618643 bp, contig 2: 1508600 bp and contig 3: 1662471 bp).

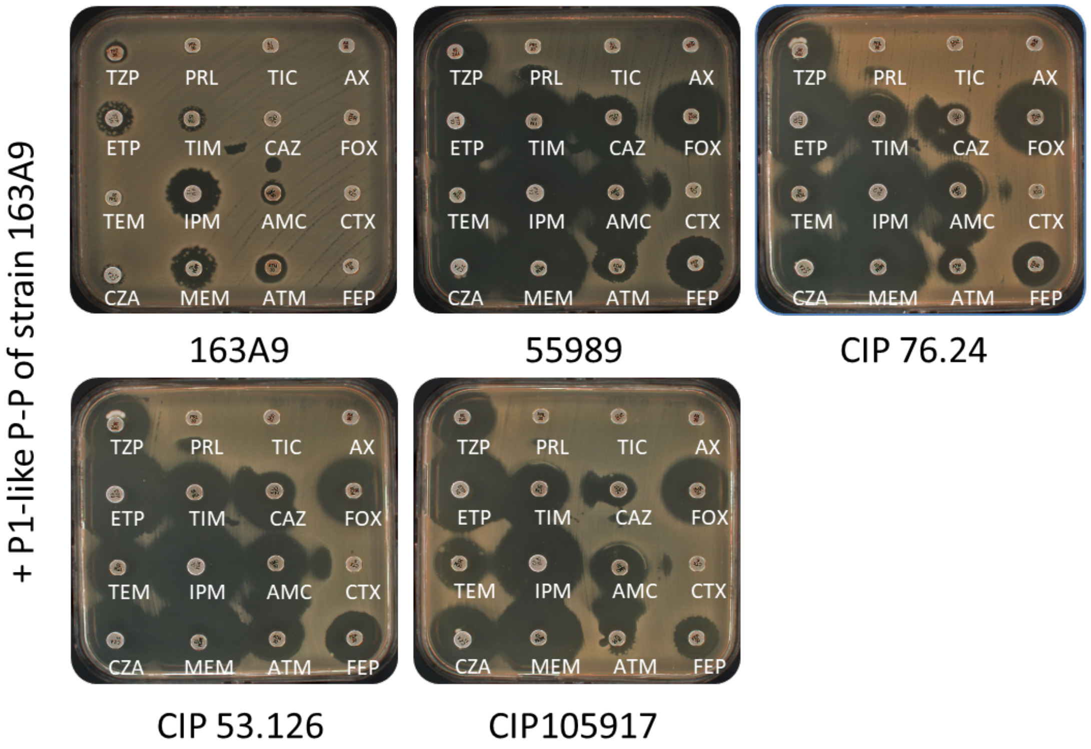

TZP: piperacillin/tazobactam
PRL: piperacillin
TIC: ticarcillin
AX: Amoxicillin
ETP: ertapenem
TIM: Ticarcillin/clavulanate
CAZ: ceftazidime
FOX: cefoxitine
TEM: temocillin
IPM: imipenem
AMC: Amoxicillin/clavulanate
CTX: cefotaxime
CZA:ceftazidime/avibactam
MEM: meropenem
ATM: aztreonam
FEP: cefepime

Figure S13: Antibiotic susceptibility test of lysogens.

*E. coli* lysogens (CIP 105197, CIP 53.126, CIP 76.24, 55989, CRE_163A9) having the P1-like P-P of strain 163A9 with the ARG *ctx-m-55* were tested for sensitivity against various β-lactam antibiotics (see Methods).

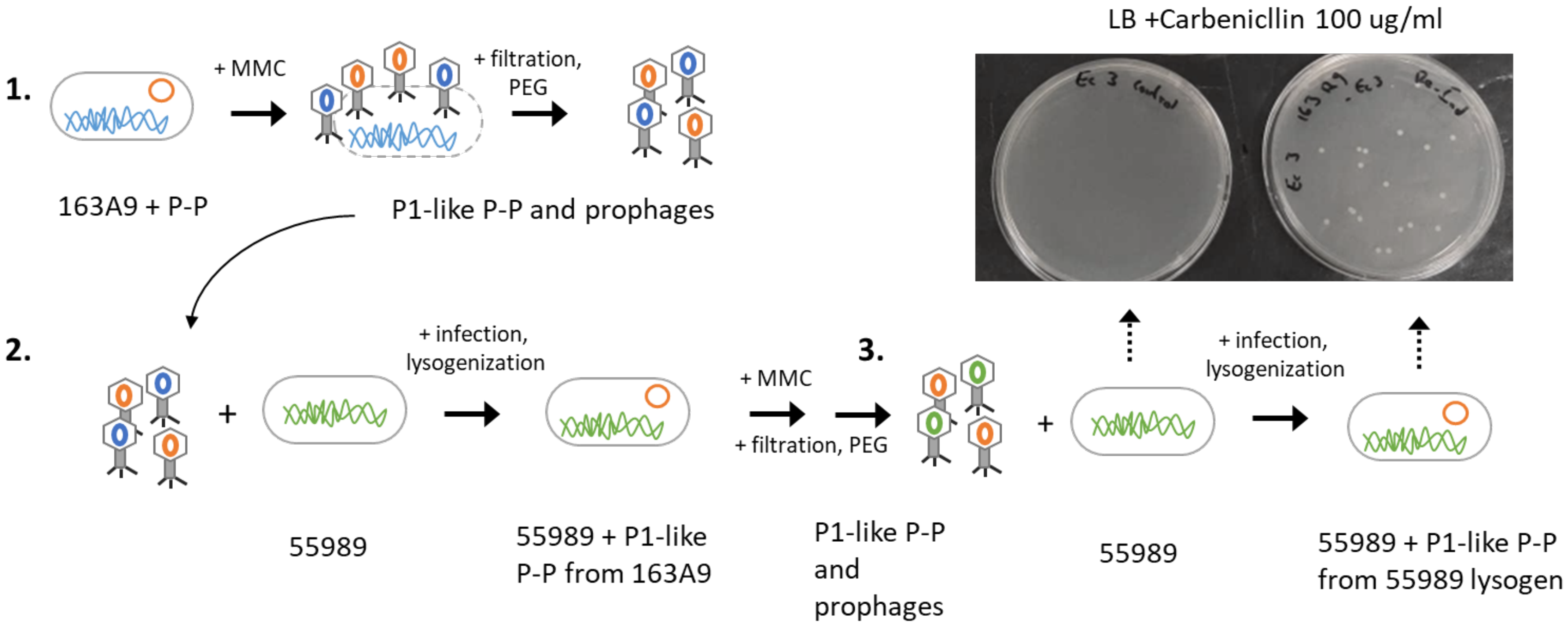

Figure S14: Re-induction and infection experiment with the P1-like P-P from 163A9
[1] *E. coli* strain 163A9 was treated with MMC (see methods) to induce the P1-like P-P. After 3-4 h, phage particles were purified and [2] subsequently used to infect *E. coli* 55989. (3) Lysogenized *E. coli* 55989 having the P1-like 163A9 P-P were exposed to MMC to induce the P-P again. Phage particles were purified, used to infect a non-lysogen variant of *E. coli* 55989 and plated on LB plates w/ 100 µg/ml Carbenicillin to screen for lysogens. In the control, un-treated *E. coli* 55989 cells were plated on LB plates w/ 100 µg/ml Carbenicillin.
